# Phase-amplitude coupling and phase synchronization between medial temporal, frontal and posterior brain regions support episodic autobiographical memory recall

**DOI:** 10.1101/2021.09.06.459104

**Authors:** Nicolas Roehri, Lucie Bréchet, Martin Seeber, Alvaro Pascual-Leone, Christoph M. Michel

**Author notes:** Corresponding author: Christoph M. Michel, Campus Biotech, Neuroscience Department, University of Geneva, 9 chemin des Mines, 1211 Genève 20, Switzerland. The first two authors contributed equally.

## Abstract

Episodic autobiographical memory (EAM) is a complex cognitive function that emerges from the coordination of specific and distant brain regions. Specific brain rhythms, namely theta and gamma oscillations and their synchronization, are thought of as putative mechanisms enabling EAM. Yet, the mechanisms of inter-regional interaction in the EAM network remain unclear in humans at the whole brain level. To investigate this, we analyzed EEG recordings of participants instructed to retrieve autobiographical episodes. EEG recordings were projected in the source space, and time-courses of atlas-based brain regions-of-interest (ROIs) were derived. Directed phase synchrony in high theta (7-10 Hz) and gamma (30-80 Hz) bands and high theta-gamma phase-amplitude coupling were computed between each pair of ROIs. Using network-based statistics, a graph-theory method, we found statistically significant networks for each investigated mechanism. In the gamma band, two sub-networks were found, one between the posterior cingulate cortex (PCC) and the medial temporal lobe (MTL) and another within the medial frontal areas. In the high theta band, we found a PCC to ventromedial prefrontal cortex (vmPFC) network. In phase-amplitude coupling, we found the high theta phase of the left MTL biasing the gamma amplitude of posterior regions and the vmPFC. Other regions of the temporal lobe and the insula were also phase biasing the vmPFC. These findings suggest that EAM, rather than emerging from a single mechanism at a single frequency, involves precise spatio-temporal signatures mapping on distinct memory processes. We propose that the MTL orchestrates activity in vmPFC and PCC via precise phase-amplitude coupling, with vmPFC and PCC interaction via high theta phase synchrony and gamma synchronization contributing to bind information within the PCC-MTL sub-network or valuate the candidate memory within the medial frontal sub-network.

## Introduction

Episodic autobiographical memory (EAM) is a complex cognitive function that involves a specific large-scale network of interacting brain regions to store and later retrieve our lives’ experiences. However, the precise mechanisms that enable the specialized brain areas to encode or retrieve EAM remain unclear. Brain oscillations and synchronization within and across large-scale networks facilitate long-distance communication and promote cognitive functions in the healthy human brain (Buzsáki 2006). Particularly, the slower theta (3-10Hz) and faster gamma (>25Hz) oscillations have been related to memory processes but likely play distinct mechanistic roles. Gamma oscillations have been suggested related to memory-related synaptic changes (Nakazono et al. 2018; Sederberg et al. 2007). Gamma phase synchrony in the neocortex is well-known to facilitate neuronal communication and functional integration thanks to the precise temporal coincidence of spiking activity (Fries 2009), and in turn binds distributed memory representations that are needed to retrieve episodic memories (Düzel, Penny, and Burgess 2010; Fell and Axmacher 2011). Theta oscillations are the dominant rhythms recorded in the hippocampus (Buzsaki 2002) and appear to serve as a mechanism of EAM memories constituted of shorter sub-episodes, whose temporal dynamics during recall are supported by the same mechanism as during encoding (Buzsaki 2002; Buzsáki and Moser 2013). Theta oscillations have been associated with both encoding and retrieval of episodic memories (Friese et al. 2013; Osipova et al. 2006; Sederberg et al. 2003).

Generally, theta phase synchrony is supposed to coordinate the coactivation of distant regions whose local neuronal assembly would be synchronized in the gamma range (Fell and Axmacher 2011). This hypothesis can also be seen as cross-frequency phase-amplitude coupling (PAC); the gamma activity would occur at a specific phase of the low-frequency oscillation (Canolty and Knight 2010; Fell and Axmacher 2011; Florin and Baillet 2015; Sirota et al. 2008; Tort et al. 2008). The low-frequency oscillation here acts as a metronome that coordinates neural activity in the same region or a remote region, making a putative mechanism for long-range communication. Using simultaneous recordings from both the hippocampus and neocortex in behaving rats and mice, Sirota et al. (Sirota et al. 2008) showed that the theta oscillations in the hippocampus modulate gamma oscillations in the neocortex through PAC. Thus, the coupling between hippocampal theta and neocortical gamma oscillations seems to be a critical mechanism supporting the large-scale memory network’s interactions (Axmacher et al. 2010; Nyhus and Curran 2010).

Very few human studies to date have explored the complex communication within and between regions involved in EAM. A few recent magnetoencephalography (MEG) studies of personal memories have started to investigate the mechanisms for the organized communication within and between broad networks at different spatiotemporal scales (Fuentemilla et al. 2014; Hebscher, Meltzer, and Gilboa 2019). For example, Fuentemilla et al. (Fuentemilla et al. 2014) found selective phase-synchronization in theta frequency between the medial temporal lobe (MTL), medial prefrontal cortex, and precuneus that was higher during EAM than general semantic retrieval. In a follow-up study, Fuentemilla et al. (2018) sought to identify larger-scale patterns in higher frequencies and found neural synchrony, specifically in the gamma frequency range underlying episodic autobiographical recollection. Hebscher et al. (2019) used theta-burst transcranial magnetic stimulation to examine whether interactions between theta and gamma frequencies facilitate the recall of personal memories and found that memory retrieval is supported by MTL-cortical communication mediated by theta phase coupling and theta-gamma PAC. Although such research emphasizes the importance of brain oscillations in memory processing and integration, these studies relied either on seed-based methods calculating the phase-synchrony between a selected region and the rest of the brain, thus missing potential interactions between regions not interacting with the seed, or scalp level connectivity analysis without any consideration of the source-level network reconstruction. Therefore, there is a need to analyze the interactions both at the source level and between all brain regions without selecting only some brain areas based on their coactivation during the EAM processes or a priori knowledge to obtain a complete picture of the interaction between brain regions.

In the present study, we investigated the dynamic interactions between neuronal networks during EAM retrieval, particularly in the gamma and high theta frequencies. We focused on the upper range of the theta band (7-10 Hz; high theta) because, while the frequency of the human hippocampal theta is rather controversial (Jacobs 2014; de la Prida 2019; Miller et al. 2018), Fuentemilla et al. (2014) found that when participants were asked to retrieve personal episodes and elaborate them during 30 s during MEG recording, the peak theta frequency linked to the MTL-Precuneus-mPFC phase synchronization was above 7 Hz. We assessed the directed phase connectivity and phase-amplitude coupling at play during EAM recollection, leveraging the high-temporal resolution of EEG and the 64 channels dataset reported in (Bréchet et al. 2019). We first computed the average power in high theta and gamma bands in the source space to investigate which parts of the brain were more active during the EAM retrieval than during a control task. We then applied two types of directed connectivity measures, the directed phase lag index (dPLI) (Stam and van Straaten 2012) and the normalized modulation index (nMI) (Özkurt and Schnitzler 2011; Tort et al. 2008) to quantify the coupling at the whole-brain level between 84 regions of interest (ROIs). The dPLI is a measure of phase-synchronization, which is robust to source leakage and analyzes the direction of the information flow. The nMI allows quantifying the PAC between the phase of low-frequency time-series and the amplitude of the high-frequency course.

## Materials and Methods

### Participants

15 healthy participants (30.5 ± 5.5 years, 10 female) were included in the study. The dataset was acquired and described in Bréchet et al. 2019. The local ethics committee’s institutional review board approved this study, and all participants provided written informed consent prior to the experiment.

### Experimental paradigm

Participants were instructed to either retrieve episodic, self-related memories (“memory condition”) or perform arithmetic calculations (“math condition”) while undergoing hd-EEG recording. There were 40 trials in total, each trial lasting 30 s. On each trial, a personal photograph (e.g., photo of a participant with a birthday cake) or a calculation (e.g., 447-7=) was presented for 3 s, followed by 27 s period of closed eyes, during which participants either retrieved the EAM or subtracted/added numbers. Participants were asked to retrieve the personal events in as much detail as possible or fully focus on the calculations. At the end of each trial, participants were asked to rate how much they were able to either relive the past self-related events or pay attention to the self-unrelated calculation. The ratings were similar between the memory and math conditions. For more detail on the experimental paradigm, see (Bréchet et al. 2019).

### EEG recording and preprocessing

EEG was recorded with a 64-channel BrainAmp EEG system (Brain Products, Munich, Germany) at 5000Hz. The EEG recording was down-sampled to 500Hz and filtered using a zero-phase digital 4th-order Butterworth bandpass filter between 1Hz and 80Hz. Infomax-based Independent Component Analysis (ICA) was applied to remove oculomotor, cardiac, and muscle artifacts based on the channels with maximal amplitude, the topography, and time course of the component (as recommended in Jung et al. 2000). Particular attention was devoted to removing long-standing muscle artifacts, which may overlap and contaminate the genuine gamma activity produced by the brain in the same frequency band (30-80Hz). In other words, we removed independent components whose topographies were limited to one or two channels and whose time-courses showed strong activity in the gamma band (Jung et al. 2000). Bad electrodes were interpolated using a 3-D spherical spline (Perrin et al. 1989) and the data were re-referenced to the common average reference. Finally, the remaining artifacts were marked and excluded from further analysis. The pre-processing stages were performed using the Fieldtrip Toolbox (Oostenveld et al. 2011).

### EEG source imaging

The forward model for source localization, based on realistic head geometry and conductivity data considering skull thickness (Locally Spherical Model with Anatomical Constraints [LSMAC]) (Brunet, Murray, and Michel 2011), was calculated for a standard set of 64-electrode positions co-registered to the Montreal Neurological Institute (MNI) atlas. The inverse solution space consisted of about 5000 point sources equally distributed in gray matter volume. We computed the inverse solution matrix using the linear distributed source localization procedure LAURA (Grave De Peralta Menendez et al. 2004). The lead-field matrix and the inverse matrix were computed using the freely available software Cartool (Brunet et al. 2011; Michel and Brunet 2019); https://sites.google.com/site/cartoolcommunity/.

### Frequency-band power analyses

We computed the analytic signal of the EEG traces using the Hilbert transform and projected the real and imaginary parts in the inverse space using the inverse operator to obtain the average power in high theta and gamma bands at each point source for the memory and math conditions. Before the Hilbert transform, the signals were bandpass filtered in either high theta (7-10Hz) or gamma (30-80Hz) bands. The square of the norm of the resulting three-dimensional complex signal was calculated to obtain a single real power time-course for each point source. A standardization across time was applied for each point source to eliminate activation biases (Bréchet et al. 2019; Michel and Brunet 2019). These values were then averaged across time and trials, resulting in an average power value at each participant’s source location.

#### Whole-brain connectivity analysis

We used 84 regions of interest (ROIs) to build the connectivity matrices. In Cartool, each source was assigned to one of the ROIs defined by the second version of the AAL atlas (Rolls, Joliot, and Tzourio-Mazoyer 2015; Tzourio-Mazoyer et al. 2002). S1 Figure shows the position of the centroid of each ROI in a 3D brain mesh and S1 Table lists the abbreviations and anatomical descriptions of the AAL ROIs.

Notably, the AAL atlas is an anatomical parcellation of the brain based on gyri and sulci rather than functional specificity. Therefore, we used several ROIs to describe functional brain areas. For example, the vmPFC was described as in Gilboa and Marlatte 2017 and McCormick et al. 2018 with the three following ROIs: the medial orbital part of the superior frontal gyrus, gyrus rectus and the subcallosal gyrus. We projected the filtered data in either high theta or gamma bands into the source space to obtain the different ROIs’ time-courses. We took the first component of the singular value decomposition computed over all the time-series of every source (independently of their orientation) belonging to a given ROI. This method was described in Rubega et al. 2019 and used in several other studies (Carboni et al. 2019, 2020; Damborská et al. 2020). The singular value decomposition has a sign ambiguity, which can bias directed connectivity measures. To resolve this indeterminacy, we used the method proposed by Bro, Acar, and Kolda 2008, which allocates the sign based on the majority of vectors represented by the component. The non-artifacts sections of the trials were concatenated, and the following connectivity measures were calculated.

We used two types of directed connectivity measures to quantify the coupling between ROIs: the directed Phase Lag Index [dPLI] (Stam and van Straaten 2012) and the normalized Modulation Index [nMI] (Canolty et al. 2006; Özkurt and Schnitzler 2011).

#### Directed Phase Lag Index

The dPLI measures the probability that the difference of the instantaneous phases of two signals is positive, i.e., the probability that one signal has its instantaneous phase leading the instantaneous phase of another signal. Formally:

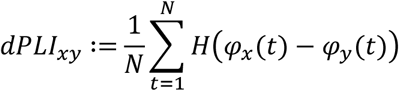

where H is the Heaviside step function, N is the number of samples, *φ*_*x*_ and *φ*_*y*_ are respectively the instantaneous phase of the time series x and y obtained after applying the Hilbert transform. *dPLI*_*xy*_ is bounded between 0 and 1. By definition, x is phase leading compared to y when *dPLI*_*xy*_ > 0.5 and phase lagging when *dPLI*_*xy*_ < 0.5. This measure was applied to the high theta and gamma filtered time-series separately.

#### Normalized Modulation Index

The nMI quantifies the phase-amplitude coupling (PAC) between the phase of low-frequency time-series and the amplitude of the high-frequency course. Mathematically:

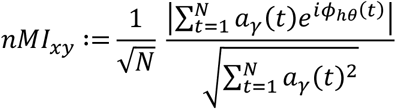

*a*_*γ*_ is the amplitude of the analytic signal filtered in gamma, *ϕ*_*hθ*_ is the instantaneous phase of the analytic signal filtered in high theta, and N is the number of samples. Note that the direction of the connectivity does not carry the usual causality meaning but rather that the phase of region A is coupled to the amplitude of region B (i.e., A →B means that 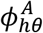 is coupled to 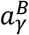).

### Statistical analysis

#### Activation maps

We applied a two-sided paired t-test to each point source’s average power across participants to investigate which parts of the brain were more active during the memory than the math condition at the group level. A cluster-based permutation procedure implemented in FieldTrip (Maris and Oostenveld 2007; Oostenveld et al. 2011) was used to correct multiple comparisons. Clusters are formed by grouping together neighboring point sources whose t-values were higher than the initial threshold. The initial threshold for cluster definition was set to p < 0.0005, and the final threshold for significance of the summed t value within clusters was set to p < 0.05 with 5000 permutations.

#### Connectivity Networks

Networks are defined by nodes and links. In this study, the nodes are the ROIs, and the links are the values of the connectivity of each pair of nodes. We used the network-based statistic (NBS) (Zalesky, Fornito, and Bullmore 2010) to compare the connectivity matrices. First, the connectivity matrices were subjected to a t-test and then corrected for multiple comparisons using a connected-component-based permutation procedure. In graph theory, a connected component is a set of nodes where any two nodes (e.g., A and B) are connected either directly (A-B) or indirectly (i.e., via other nodes: A-C-D-B). As for the cluster-based statistics, one can derive a measure for each component present after the first level threshold, presently the number of links, and test this measure against values obtained after permutation. Importantly, NBS only works for undirected networks as it uses a breadth-first search (Hopcroft and Tarjan 1973). We modified this part to work with a directed network by searching for weakly connected components, i.e., seeking connected components while ignoring link direction as implemented in Matlab’s *conncomp*. We chose weakly over strongly connected components because strongly connected components have a stricter definition, which may not be realistic in terms of brain networks. A component is defined as strongly connected if and only if for every pair of nodes, there is a directed path between any two nodes. For instance, A→B→C is weakly connected because there is no directed path to go from B or C to A, while A→B→C→A is strongly connected, as there is a directed path that connects B to A (B→C→A) and C to A (C→A).

As proposed by Zalesky and colleagues (Zalesky et al. 2010), the data can be transformed prior to the t-test to ensure a meaningful first level threshold. Both the nMI and dPLI are bounded by 0 and 1. Therefore we applied a logit transform. The logit transform is a monotonically increasing function and transforms values from(0,1) to(–∞,+∞), specifically logit(0.5) = 0. The above inequalities relative to the dPLI are conserved except that the critical value became 0 instead of 0.5.

We directly contrasted the memory and math conditions through a paired t-test as the nMI matrices are non-symmetric (*nMI*_*xy*_ ≠ *nMI*_*yx*_). The permutations were obtained by randomly exchanging the conditions within subjects. However, the dPLI matrices are anti-symmetric (*dPLI*_*xy*_= –*dPLI*_*yx*_) and a direct comparison of the conditions is thus not possible. Therefore, we first applied a one-sample t-test against 0 to determine the presence of a preferred direction between nodes across participants and then removed the links present in both memory and math conditions. Here the permutations were obtained by randomly transposing the matrices, i.e., changing the direction of the links. The initial threshold for connected-component definition was set to p < 0.005. The final threshold for significance of the number of connections within connected components was set to p < 0.05 with 5000 permutations. We also defined three different types of nodes: a.) source nodes are nodes with only outgoing links, b.) sink nodes with only incoming links, and c.) intermediate nodes have incoming and outgoing links. When describing the significant networks, we will give their number of links and nodes and, for the sake of readability, only detail the connectivity of their strongest nodes, i.e., nodes with more than three links. The complete networks are shown in the respective figures and supplementary tables.

#### Comparison of the distance between connected regions

Finally, we calculated the distance between the centroids of pairs of connected ROIs for each connectivity type. We performed a one-way ANOVA to test whether there exists a difference in these profiles of distance. Post hoc t-tests were used to explore differences between all the pairs of connectivity types and were controlled for multiple comparisons by the Dunn-Sidák approach (Šidák 1967). All reported p-values were corrected for multiple comparisons by one of the above approaches.

## Results

### Power changes in source space

We found several statistically significant clusters of point sources in gamma and high theta bands when comparing the power in the memory condition to the control/math condition **(Fig. 1, Table 1)**. Specific to the gamma frequency, we found 5 significant clusters (Fig. 1a, Table 1a). The first cluster (sum of t: 982.03, p < 0.01) had its maximum t-value located in the right superior temporal gyrus (STG), and the right temporal pole and supramarginal gyrus (SMG) were its anterior and posterior limits, respectively. It also included part of the right middle temporal gyrus (MTG), insula, rolandic operculum, and postcentral gyrus. The second cluster (sum of t: 624.44, p < 0.01) showed a maximum t-value in the left precuneus (PrCu) and included the PrCu bilaterally and the left superior parietal (SPG) and inferior parietal gyri (IPG). The third cluster (sum of t: 258.80, p < 0.01) bilaterally included the median cingulate cortex (MCC) and posterior superior frontal gyrus (pSFG), and its peak was in the left MCC. The peak of the fourth cluster was found in the left SMG, and this cluster (sum of t: 89.88, p<0.01) was composed of the left SMG, IPG, and postcentral gyrus. The fifth cluster (sum of t: 32.29, p<0.05) included the right caudate nucleus and thalamus with a peak in the right caudate nucleus.

**Table 1.** Anatomical description of the statistically significant clusters obtained when contrasting memory and math condition in the a. gamma band and b. high theta band.

| **a. gamma band** |  |  |  |  |  |  |  |  |
| --- | --- | --- | --- | --- | --- | --- | --- | --- |
| **# Cluster** | **size** | **p** | **AAL regions** | **peak region** | **peak t** | **X** | **Y** | **Z** |
| 1 | 193 | 0.0014 | R STG<br>R MTG<br>R insula<br>R rolandic operculum<br>R PostCG<br>R Putamen<br>R SMG<br>R STGpole<br>R MTGpole<br>R heschl gyrus<br>R ITG<br>R Pallidum | R STG | 6.51 | 55 | -3 | -3 |
| 2 | 128 | 0.0032 | L precuneus<br>R precuneus<br>L SPG<br>L IPG<br>L SOG<br>L MCC<br>L cuneus<br>R MCC<br>L MOG | L precuneus | 5.68 | -3 | -55 | 42 |
| 3 | 54 | 0.0056 | L MCC<br>R MCC<br>L pSFG<br>R pSFG | L MCC | 5.29 | -3 | -10 | 42 |
| 4 | 19 | 0.0090 | L SMG<br>L IPG<br>L PostCG | L SMG | 5.27 | -62 | -29 | 36 |
| 5 | 7 | 0.0130 | R Caudate<br>R Thalamus | R Caudate | 4.70 | 16 | -3 | 23 |
| **b. high theta band** |  |  |  |  |  |  |  |  |
| **# Cluster** | **size** | **p** | **AAL regions** | **peak region** | **peak t** | **X** | **Y** | **Z** |
| 1 | 37 | 0.0030 | L SPG<br>L PostCG<br>L IPG<br>L precuneus<br>L PreCG | L SPG | 7.10 | -36 | -62 | 62 |
| 2 | 13 | 0.0078 | R SFG<br>R pSFG | R SFG | 4.89 | 29 | -3 | 68 |
| 3 | 12 | 0.0080 | L pSFG<br>R pSFG<br>L SFG | L pSFG | 5.05 | -10 | -3 | 74 |
| 4 | 4 | 0.0120 | R SPG<br>R PostCG | R SPG | 5.12 | 29 | -55 | 68 |

*The lists of AAL regions included in a cluster are sorted in a descending order according to their number* *of point sources within the cluster. Sum of t-values correspond to the summation of the t-values within*
a cluster. The size corresponds to the number of point sources making up the cluster. The region where the peak t-value is found is given under the peak region column. X, Y and Z are MNI coordinates in mm of the peak. Only regions with more than 1% of their size included in the clusters are listed. Annotations: L: left, R: Right, STG: superior temporal gyrus, MTG: middle temporal gyrus, ITG: inferior temporal gyrus, PreCG: precentral gyrus, PostCG: postcentral gyrus, SMG: supramarginal gyrus, SPG: superior parietal gyrus, IPG: inferior parietal gyrus, SOG: superior occipital gyrus, MOG: middle occipital gyrus, MCC: median cingulate cortex, SFG: superior frontal gyrus, pSFG: posterior superior frontal gyrus.

**Fig. 1.**
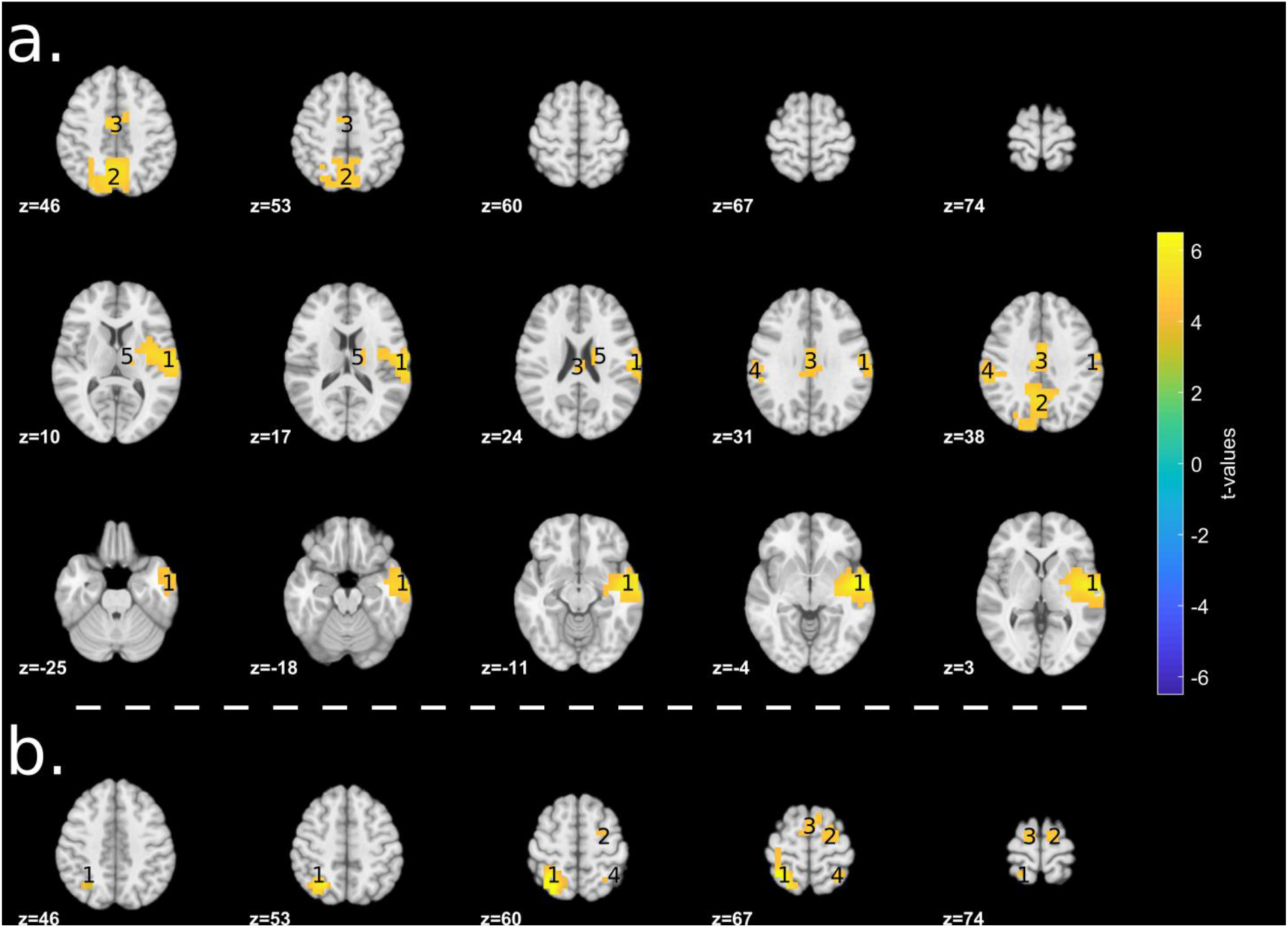
Significant clusters in a. gamma and b. high theta band when contrasting the memory vs. math conditions. Only slices containing significant clusters are shown. Numbers identify the different clusters and are detailed in Table 1

We identified 4 significant clusters in the high theta frequency (Fig. 1b, Table 1b). The first cluster (sum of t: 204.54, p < 0.01) had its maximum t-value located in the left SPG and included the left postcentral gyrus, IPG, precentral gyrus, and the left PrCu. The second cluster (sum of t: 60.62, p < 0.01) showed a peak in the right superior frontal gyrus (SFG) and included the right pSFG. The third cluster (sum of t: 56.78, p < 0.01) bilaterally included part of the pSFG and the left SFG with a peak in the left pSFG. The fourth cluster (sum of t: 18.87, p < 0.05) showed a maximum t-value in the right SPG and partially included the postcentral gyrus.

### Directed functional connectivity

As shown in **Figure 2 and S2 Table**, the gamma band directed connectivity analysis revealed a network (p < 0.01) composed of 16 links and 16 nodes (including 9 source, 1 intermediate, and 6 sink nodes). No statistically significant network was found in the math condition, and therefore no link was removed in the network found in the memory condition as described in the Method section. Most of the links connected nodes in the frontoparietal midline and left MTL. Particularly, the amygdala (AMY) was the sink node with the most incoming links (4 links) originating from the hippocampus (HIPP), parahippocampal gyrus (PHG), thalamus (Th), and left PCC. The right subcallosal gyrus (SCG) was the second strongest sink node with 3 links connected to the right PCC, pSFG, and left Th. With the same number of incoming links, the left anterior cingulate cortex (ACC) was connected to the left and right pSFG and the medial orbital part of the right SFG (SFGmedOrb). The pSFG was the source node with the most outgoing links, connecting the ACC and right SCG.

**Fig. 2.**
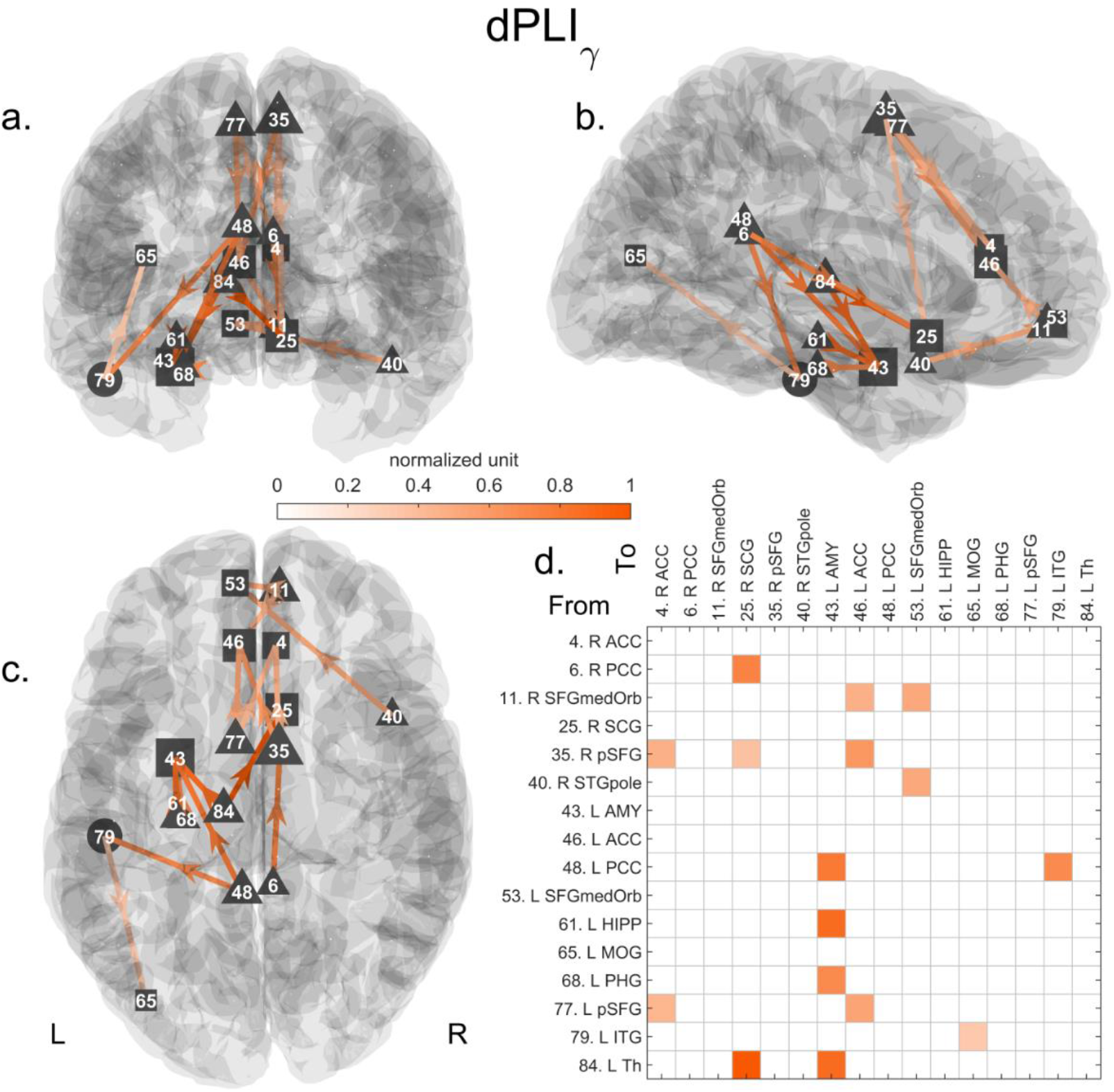
Significant directed network in gamma in the memory condition. The statistically significant network (16 links and 16 nodes, p<0.01) is represented in 3D along the a.) coronal, b.) sagittal and c.) axial axes. The connectivity matrix is shown in d. Triangles and squares encode nodes with only outgoing or incoming links, i.e., source or sink nodes, respectively. Circles correspond to intermediate nodes with both incoming and outgoing links. The normalized grand average of the connectivity is color-coded. One corresponds to the maximum value. The size of the node encodes its number of (outgoing or incoming) links. Annotations: L: left, R: Right, Th: thalamus, SCG: subcallosal gyrus, AMY: amygdala, HIPP: hippocampus, PCC: posterior cingulate cortex, ITG: inferior temporal gyrus, PHG: parahippocampal gyrus, SFG: superior frontal gyrus, pSFG: posterior SFG, ACC: anterior cingulate cortex, STGpole: polar part of the superior temporal gyrus, SFGmedOrb: medial orbital part of the SFG, ITG: inferior temporal gyrus, MOG: middle occipital gyrus.

As shown in **Figure 3 and S3 Table**, the high theta directed connectivity analysis revealed the presence of a medial posterior-to-anterior network (p < 0.01) composed of 21 links and 18 nodes (including 4 source and 14 sink nodes). Only one link was present in both memory and math conditions and was removed only to keep links specific to the memory condition as described in the Methods section. The left PCC was the source region with the most outgoing links (9 links). This region drove the right ACC and bilaterally the gyrus rectus (Rec), SCG, Th, and MCC. The right PCC was driving 5 regions: the right Rec and bilaterally the Th and SCG. The right inferior temporal gyrus (ITG) led the right Rec, SFGmedOrb, opercular part of the IFG, and the orbital part of both SFG. Interestingly, the two strongest source nodes (left and right PCCs) were mostly connected to the nodes of the vmPFC (defined by the SFGmedOrb, Rec, and SCG nodes).

**Fig. 3.**
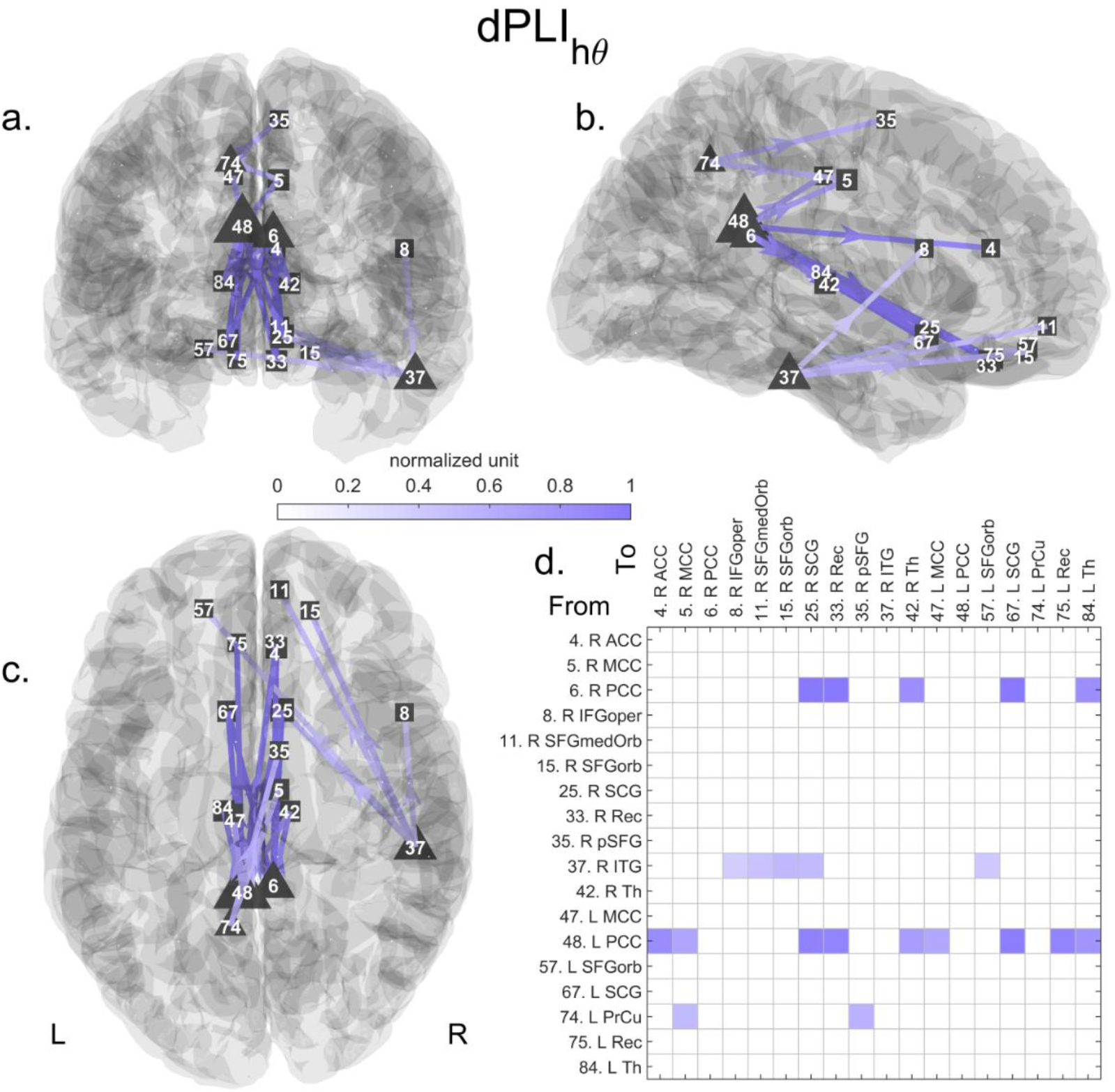
Significant directed network in high theta in the memory condition. The statistically significant network (21 links and 18 nodes, p<0.01) is represented in 3D along the a.) coronal, b.) sagittal and c.) axial axes. The connectivity matrix is shown in d. Triangles and squares encode nodes with only outgoing or incoming links, i.e., source or sink nodes, respectively. The normalized grand average of the connectivity is color-coded; one corresponds to the maximum value. The size of the node encodes its number of (outgoing or incoming) links. Annotations: L: left, R: Right, PCC: posterior cingulate cortex, SCG: subcallosal gyrus, Rec: gyrus rectus, ACC: anterior cingulate cortex, Th: thalamus, MCC: median cingulate cortex, PrCu: precuneus, SFG: superior frontal gyrus, pSFG: posterior SFG, ITG: inferior temporal gyrus, SFGorb: orbital part of the SFG, ITG: inferior temporal gyrus, SFGmedOrb: medial orbital part of the SFG.

### Phase-amplitude coupled network

Compared to the gamma and high theta directed networks, the direction of the PAC-directed network links carries information about the phase to amplitude relationship. For a given pair of connected nodes, the high theta phase of the node with the outgoing link is coupled to the gamma amplitude of the node with the incoming link. We identified a statistically significant PAC network (p < 0.05) when comparing the memory to the math conditions **(Fig. 4 and S4 Table)**. This network was composed of 20 links and 20 nodes (including 12 source, 3 intermediate, and 5 sink nodes). The left SFGmedOrb (part of the vmPFC) was the sink region with the most incoming links (5 links). Its gamma amplitude was coupled to the high theta phase of the left PHG, HIPP, STG, ITG, and lingual gyrus. The right SPG was the second sink node with the most incoming links (4 links). It was coupled to the left HIPP and MCC, and right AMY and superior occipital gyrus. The right Rec (also a node of the vmPFC) had 3 links coming from the right MTG pole, right insula, and left MTG. The left fusiform gyrus received 3 connections from the right STG, precentral gyrus, and angular gyrus. Interestingly, we found that the left HIPP had its high theta phase coupled to gamma amplitude of the two sink nodes with the highest number of links, namely the SFGmedOrb (part of the vmPFC) and right SPG.

**Fig. 4.**
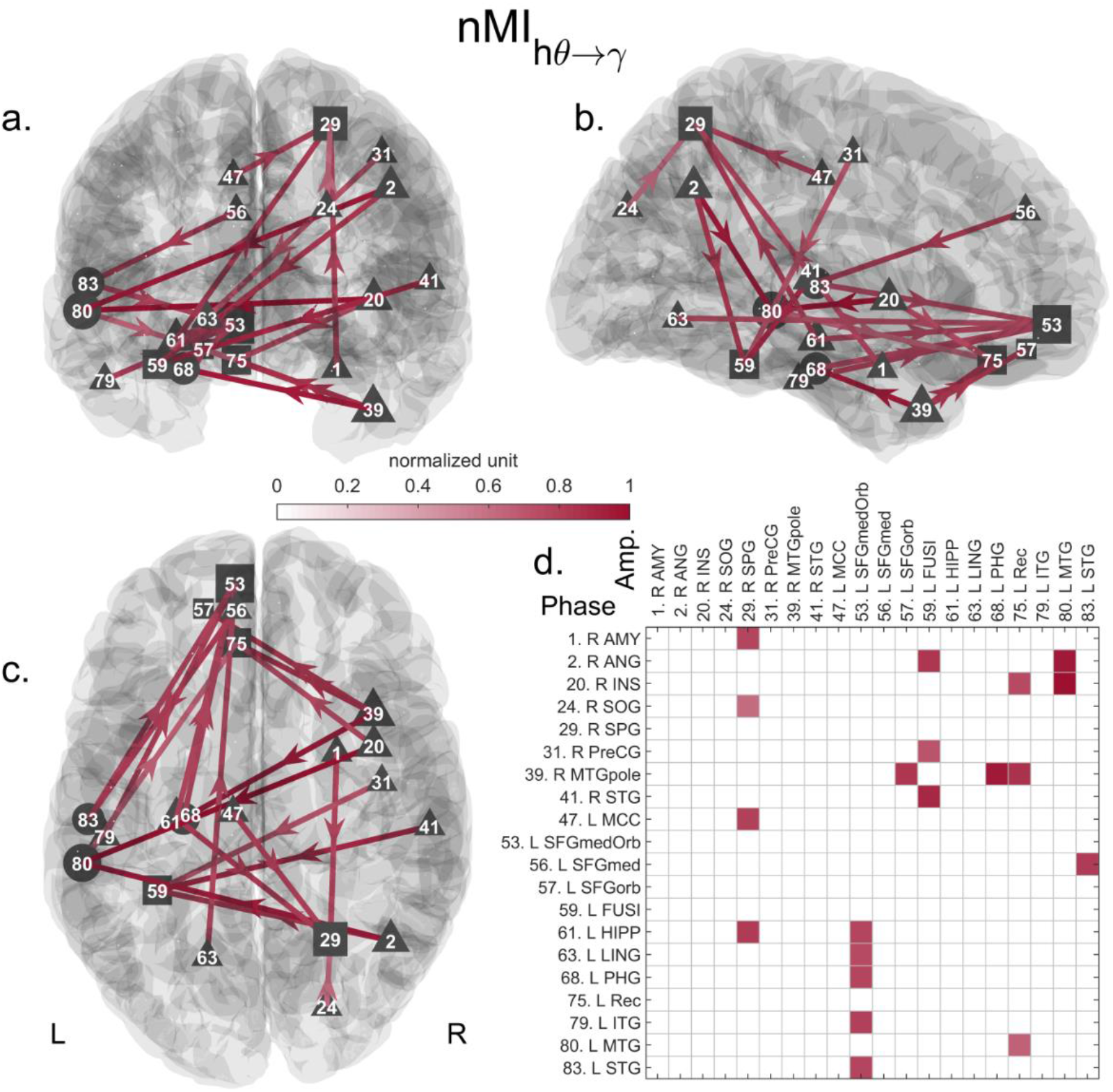
Significant phase-amplitude coupling directed network when contrasting the memory and math conditions. The statistically significant network (20 links and 20 nodes, p<0.05) is represented in 3D along the a.) coronal, b.) sagittal and c.) axial axes. The connectivity matrix is shown in d. Triangles and squares encode nodes with only outgoing or incoming links, i.e., source or sink nodes, respectively. Circles correspond to intermediate nodes with both incoming and outgoing links. The normalized grand average of the connectivity is color-coded. One corresponds to the maximum value. The size of the node encodes its number of (outgoing or incoming) links. Annotations: L: left, R: Right, INS: insula, MTG: middle temporal gyrus, ANG: angular gyrus, MTGpole: polar part of the MTG, PHG: parahippocampal gyrus, STG: superior temporal gyrus, FUSI: fusiform gyrus, Rec: gyrus rectus, SFG: superior frontal gyrus, SFGorb: orbital part of the SFG, HIPP: hippocampus, SPG: superior parietal gyrus, SFGmed: medial part of the superior frontal gyrus, MCC: median cingulate cortex, ITG: inferior temporal gyrus, SFGmedOrb: medial orbital part of the SFG, AMY: amygdala, LING: lingual gyrus, PreCG: precentral cortex, SOG: superior occipital gyrus.

### ROI to ROI distance profile in different connectivity type

We applied one-way ANOVA to test whether the distributions of the distances between two linked ROIs were different across connectivity types and found a significant result (F = 11.32, p < 0.0001, Fig. 5). Furthermore, the post-hoc t-test revealed that the distances were higher when the ROIs were connected by PAC than by high theta (p < 0.05) or gamma phase synchrony (p < 0.001).

**Fig. 5.**
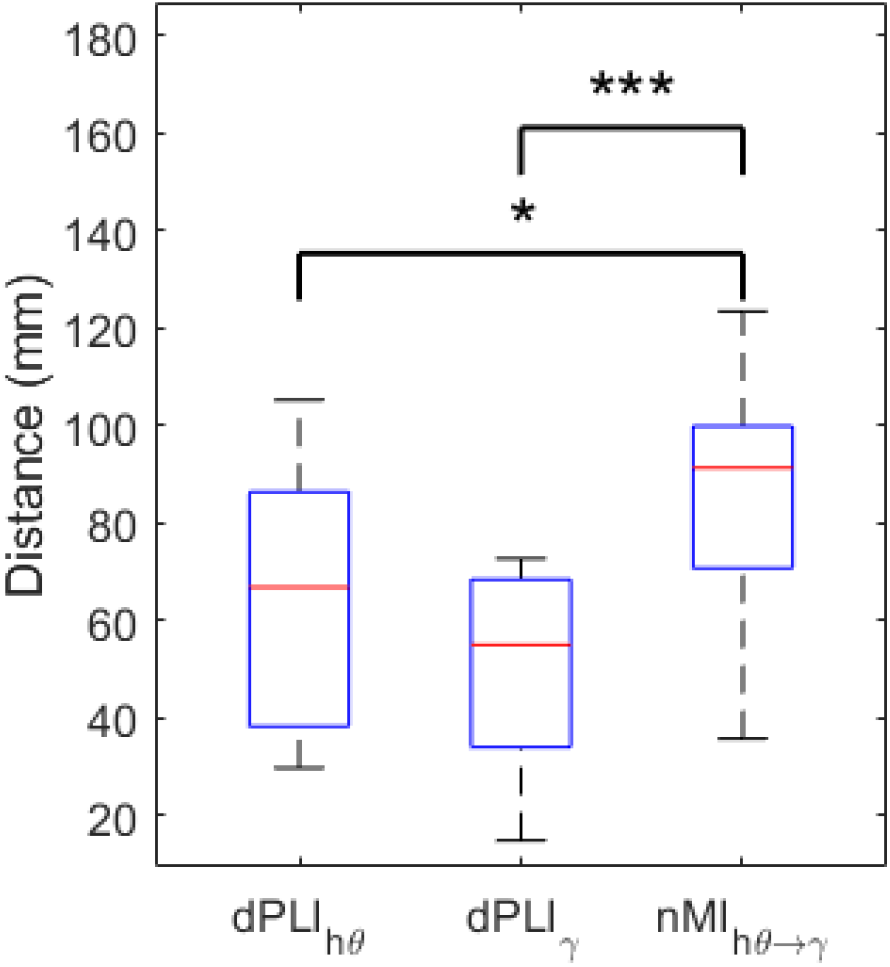
Boxplot of the distances between ROIs connected either by high theta, gamma phase synchrony, or phase-amplitude coupling. The one-way ANOVA showed that these distributions were not originating from the same distribution (F = 11.32, p < 0.0001). Post-hoc t-tests indicated that the distances between ROIs connected via phase-amplitude coupling (nMI) were longer than via phase synchrony. (*: p<0.05, ***: p<0.001, the reported p-value were adjusted after the Dunn-Sidák procedure correcting for multiple comparisons). dPLI: directed phase lag index, nMI: normalized Modulation Index, hθ: high theta, γ: gamma.

## Discussion

### A brief summary of main findings

Our results showed frequency-specific activation of PrCu, SMG, MCC, and STG in the gamma band and SPG, SFG, and pSFG in the high theta band. We also found a frequency-specific directed phase-synchrony network that highlighted the importance of frontoparietal midline and MTL interactions. Specifically, in the gamma band, the left AMY ROI was driven by other MTL regions (HIPP, PHG) and left PCC, and the pSFG, ACC, and vmPFC were interconnected. In the high theta band, a posterior-to-anterior network was found with the PCC driving the vmPFC. Finally, the high theta-gamma PAC network was dense and showed that the MTL phase organized the left vmPFC and right SPG. Overall, the results reveal distinct spatio-temporal signatures in the high theta and gamma bands, with specific interactions that we hypothesize reflect distinct cognitive processes of self-referential autobiographical memory retrieval.

### Increased gamma activity in regions linked to self-relevant processing

The brain regions with increased gamma activity were primarily found in the left PrCu, left SMG, right STG, and MCC. Generally, these gamma activity clusters seem to occur in regions whose role is related to the first-person perspective and self-orientation.

The PrCu and the SMG have been implicated in self-relevant processing. The PrCu is supposed to shift the frame of reference from an allocentric to an egocentric one (Hebscher, Levine, and Gilboa 2018), and stimulation of this region has been shown to alter the autobiographical memory retrieval (Hebscher, Ibrahim, and Gilboa 2020; Hebscher et al. 2019). In a case report, it was reported that epileptic seizures involving the left SMG provoked autonomic hallucinations (i.e., seeing the double of oneself from an internal point of view) in a patient investigated with intracerebral EEG (Fonti et al. 2020). In addition, these two regions (PrCu and SMG) were also found in the fMRI analysis in the original paper of the analyzed dataset (Bréchet et al. 2019), strengthening the implication of these regions in EAM.

The MCC is known to be involved in the head and body’s reflexive orientation in space (Vogt 2016). This region could work in concert with the PrCu and SMG areas to create an internal representation and sense of self-orientation during EAM recall. The right STG is often listed as activated during recollection of EAM (Boccia, Teghil, and Guariglia 2019; Bréchet et al. 2018) but does not belong to the core regions, unlike the anterior temporal area (Svoboda, McKinnon, and Levine 2006). Of note, it could be possible that, because of the lower spatial resolution of the EEG, the activation of the right temporal junction and the right anterior temporal area merged into a single cluster.

### Activation of the dorsal Frontoparietal Network in high theta band

Increased activation of the SFG, pSFG, and SPG was found in a high theta band in the memory condition compared to the math condition. Because the clusters found in the SFG do not entirely delineate this gyrus, it can be further interpreted as belonging to the superior frontal sulcus. These regions are known to be part of the dorsal Frontoparietal Network (dFPN) (Petersen and Posner 2012; Ptak, Schnider, and Fellrath 2017). Its functions are diverse and include motor planning and imagery, mental rotation, spatial attention, and working memory (Ptak et al. 2017). At first, it could seem unexpected to find such a visual attentional network more active in the EAM condition than in the math condition. However, during the memory condition, the participants were asked to relive past events depicted on self-related photographs for more than twenty seconds. This requires manipulating and maintaining the scene internally, which is in line with functions attributed to the dFPN. Although the math condition is a working memory task, the regions of the dFPN were less involved in that condition. Solving simple arithmetic operations rely more on the frontal network (Ishii et al. 2014). Our previous study showed that one specific microstate occurred more often during the math than memory condition (Bréchet et al. 2019). This microstate’s source localization revealed activity in the dorsal ACC, IFG, and IPG, attributed to the executive control system (Ferris and Peterson 2012).

### MTL organizes local gamma synchronization in prefrontal and parietal cortices

Our PAC result showed that the left hippocampus’ high theta phase orchestrated the gamma activity in the medial orbital part of the superior frontal gyrus (belonging to vmPFC) and right superior parietal gyrus. According to the “reader-initiated” mechanism hypothesis (Buzśaki and Wang 2012; Sirota et al. 2008), the hippocampus through theta oscillations can initiate long-range reciprocal communication with neocortical gamma oscillations. In that case, the hippocampus would act as a metronome that coordinates neocortical gamma oscillators. Those gamma oscillations then locally, i.e., within the same region, synchronize to combine information sent back to the hippocampus. We determine that the left AMY node received information from the left PCC, Th, PHG, and HIPP via gamma phase synchrony. Because of the lower spatial resolution of EEG, it might be possible that the AMY node activity globally represents the activity of the anterior MTL rather than activity solely of the AMY. Similarly, we interpret the PHG and HIPP nodes as representing the MTL activity and only refer to their name to describe their connections within the network. This gamma phase synchrony could be the reciprocal connection initiated via PAC by the left MTL with the right superior parietal gyrus, which sends information back via the PCC to the anterior MTL.

A possible role of this reciprocal connection is to perform the high-resolution contextual binding to generate a coherent scene (Yonelinas 2013). Indeed the nodes of this gamma sub-network (AMY, HIPP, PHG, PCC) form the scene-reconstruction network (Andrews-Hanna et al. 2010; Axelrod, Rees, and Bar 2017; Hassabis and Maguire 2007), which was also found in the original paper’s BOLD connectivity analysis and EEG microstate analysis (Bréchet et al. 2019). Furthermore, as defined in AAL, the PCC ROI contains part of the retrosplenial cortex (Rolls et al. 2015). This region works in concert with the Th, which sends head direction information, to switch between allocentric and egocentric perspective (Vann, Aggleton, and Maguire 2009). The anterior hippocampus, which is more engaged during scene and events reconstruction than the posterior hippocampus (Zeidman and Maguire 2016), reconstructs the scene from an allocentric viewpoint. This gamma connectivity from the PCC/retrosplenial cortex, Th, and PHG to anterior MTL could help orient EAM recollection to a personal view.

The second node entrained by the hippocampal phase was in the vmPFC. This node was part of a more extensive gamma phase synchrony sub-network constituted by the ACC, pSFG, and other nodes of the vmPFC. Fuentemilla et al. (2018) showed that large-scale gamma synchrony supports autobiographical recollection and that this synchrony also occurred in the midline frontal MEG sensor, in line with our findings. The putative role of the vmPFC is to monitor memory veracity and correctness (Barry and Maguire 2019; Eichenbaum 2017; Gilboa and Moscovitch 2017; Hebscher and Gilboa 2016; Ritchey and Cooper 2020). This frontal gamma phase synchronization could be the mechanism by which the vmPFC combines information from the prefrontal cortex to verify the retrieved memory’s validity.

Finally, when analyzing the distance between the pairs of coupled ROIs, we showed that the distance between ROIs connected via PAC was higher than that via high theta or gamma phase synchronization. PAC thus seems to be the preferred mechanism for long-range coupling, while high theta or gamma phase synchronization appears to be the preferred mechanism for local coupling. Our results suggest that the MTL by entraining with its high theta phase local gamma synchronization in remote and functionally diverse regions, namely the medial prefrontal and the posterior associative cortex, permits to reconstruct a coherent and valid self-relevant episode.

### PCC leads and vmPFC follows the network dynamics

On the one hand, we found nodes within the vmPFC as sink nodes in the high theta phase synchrony network and the PAC network. This result suggests that the vmPFC is driven via the high theta oscillations in the former network and that gamma activity is coupled to other regions’ high theta phase in the latter network. On the other hand, we found the PCC to exclusively contain source nodes in high theta and gamma phase synchrony network, i.e., the PCC was phase leading other regions. Specifically, the network analysis revealed a PCC to vmPFC connection in the high theta band. This result is supported by anatomical connections, namely the cingulum bundle, between these two regions (Bubb, Metzler-Baddeley, and Aggleton 2018) and a recent paper showing the same link in the alpha band within the Default Mode Network (Wang et al. 2019). The PCC is supposed to play a critical role in integrating relevant episodic information emerging from posterior cortices (Andrews-Hanna, Saxe, and Yarkoni 2014; Bird et al. 2015). This directed connectivity suggests that once episodic elements are integrated within the PCC, they are sectioned within high theta cycles and sequentially made available via high theta phase synchrony to the vmPFC to either maintain or withhold a memory candidate.

Furthermore, the vmPFC’s gamma activity was coupled to regions other than the left MTL (as discussed above), including the left STG, ITG, the right MTG pole, and the right insula. These regions are linked to either semantic memory processing or emotion processing and are known to contribute to EAM recollection by adding semantic and emotional features (Svoboda et al. 2006). This PAC relationship suggests that the vmPFC is phase-biased by these regions to constrain the memory validation procedure by the semantic features supporting the EAM.

The fact that the vmPFC was mainly a sink node seems to contradict the results of McCormick et al. (2020). While the memory and baseline conditions are relatively similar, our studies differ in several ways. First, they focused on the five first seconds of the trial while we analyzed the whole trial. Therefore, their finding could describe the construction phase of the EAM while ours encompasses both the construction and elaboration of EAM. Moreover, the hippocampal and vmPFC ROIs in McCormick et al. (2020) were selected by contrasting the memory and math/baseline condition and corresponded to areas with attenuation or decrease of broadband (1-30 Hz) activity. Our ROIs were based on an anatomical atlas. Finally, in McCormick et al. (2020), dynamic causal modelling was used to determine which of the hippocampus or the vmPFC was driving the other regions, which, as far as we know, does not model phase-amplitude coupling or phase synchronization (i.e., the two mechanisms considered in our investigation). Rather than contradicting each other, our study expands the findings of McCormick et al. (2020) and conveys different, complementary information about the dynamic procedure of EAM recall.

### Potential limitations and the curse of double-dipping

A recent paper from the OHBM COBIDAS committee (Pernet et al. 2020) emphasized the problem of double-dipping (Kriegeskorte et al. 2009, 2010). This problem could arise when secondary statistical analysis (in our case, the connectivity analysis) is applied to ROIs selected with the same statistical criterion using the same dataset. To avoid this issue, we could have selected ROIs with large power in both conditions or with larger power in the memory vs. math condition but using an independent dataset. Rather than selecting ROIs, we avoided double-dipping by applying whole-brain analyses. Interestingly, the regions with high theta or gamma power increase were almost not present in the three obtained networks. The PrCu, SMG, and MCC showed a greater gamma power in the memory than in the math condition, but they were absent in the gamma phase synchrony and PAC network. Likewise, the SPG and SFG were absent in the high theta phase network while showing a high theta power increase. The results would have been strikingly different if only the activated ROIs had been selected for the network analysis. In fact, ROIs with higher activity are not necessarily connected to each other as their activity could evolve independently of each other, especially in our case with trials lasting several tens of seconds. Moreover, brain areas that do not exhibit a difference in power between the conditions could be differently connected in the two conditions. These interactions would be missed if connectivity analysis is applied only to ROIs with larger activity in the condition of interest. We thus believed that applying whole-brain connectivity analysis, as proposed here, is more appropriate to capture networks completely.

PAC integrates the gamma band’s amplitude and thus could be biased by changes in amplitude. However, the regions whose gamma activity is synchronized to the high theta phase of another region are not present in the gamma’s activation map, especially the vmPFC. Therefore, the PAC network cannot be uniquely attributed to differences in gamma power between the conditions.

Finally, gamma activity in scalp EEG can be highly influenced by noise or muscle artifacts. If such artifacts were still present after ICA denoising, they would generate zero-lag connectivity between regions affected by the artefact. As the dPLI is insensitive to zero-lag connectivity and no self-loop, i.e., a region is connected to itself, is present in the PAC network, it is unlikely that artifact bias the obtained networks.

## Conclusion

In sum, our findings suggest that the left medial temporal lobe (Hipp, AMG and PHG), the ventromedial prefrontal cortex (vmPFC), and parietal regions (PCC and SPG) work in tandem during recall of self-relevant episodic memories. Importantly, this synergy occurs thanks to complex coordination via frequency-specific and cross-frequency coupling. The medial temporal lobe orchestrates the gamma activity in the ventromedial prefrontal cortex and parietal regions with its high theta phase. These two areas synchronize in phase their gamma activity with neighboring regions to, possibly, valuate and construct the memory. Finally, a high theta synchronization occurs between the posterior cingulate cortex and medial prefrontal regions, which could show the transfer of sub-episodes of memory from posterior to frontal regions. Future studies should investigate these relationships utilizing a higher number of electrodes and a paradigm allowing the separation of the activity of regions related to semantic memory to refine these episodic autobiographical memory networks further.

## Abbreviations

EAM: episodic autobiographical memory
PAC: phase-amplitude coupling
ROI: region of interest
dPLI: directed Phase Lag Index
nMI: normalized Modulation Index
NBS: network-based statistic
dFPN: dorsal Frontoparietal Network
MTL: medial temporal lobe
ACC: anterior cingulate cortex
MCC: median cingulate cortex
PCC: posterior cingulate cortex
SFG: superior frontal gyrus
SFGmedOrb: medial orbital part of the SFG
pSFG: posterior superior frontal gyrus
vmPFC: ventromedial prefrontal cortex
SCG: subcallosal gyrus
Rec: gyrus rectus
STG: superior temporal gyrus
MTG: middle temporal gyrus
ITG: inferior temporal gyrus
PrCu: precuneus
SPG: superior parietal gyrus
IPG: inferior parietal gyrus
SMG: supramarginal gyrus
PHG: parahippocampal gyrus
HIPP: hippocampus
AMY: amygdala
Th: Thalamus

## Data and Code availability statement

The participants informed consent form did not include permission to allow data to be uploaded to an online repository. Data and materials are available upon direct request and are subject to anonymization to protect the privacy of participants. Requestors must sign a formal data sharing agreement.

Data were analyzed using open source and open access tools. EEG recording were analyzed using Fieldtrip (Oostenveld et al. 2011), available at: https://github.com/fieldtrip/fieldtrip. The lead-field matrix and the inverse matrix were computed using the freely available software Cartool (Brunet et al. 2011; Michel and Brunet 2019); available at https://sites.google.com/site/cartoolcommunity/. The Network Based Statistic toolbox(Zalesky et al. 2010) is available at https://www.nitrc.org/projects/nbs.

## Funding

This work was supported by the Swiss National Science Foundation to LB (grant No. 187949) and CM (grant No. 320030_184677), and by the National Center of Competence in Research (NCCR) ‘SYNAPSY - The Synaptic Bases of Mental Diseases’ financed by the Swiss National Science Foundation (SNF, grant n° 51AU40_125759).

## Notes

### Competing Interest Statement

The authors have declared no competing interest.

