## Supplementary material for "Phase-amplitude coupling and phase synchronization between medial temporal, frontal and posterior brain regions support episodic autobiographical memory recall": S1 Figure

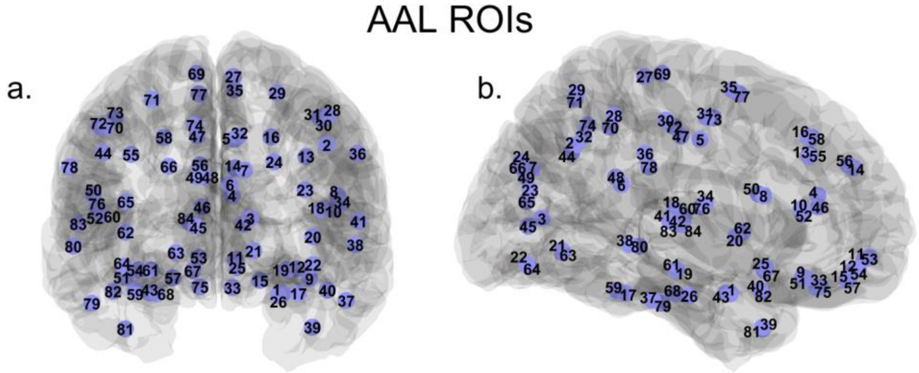


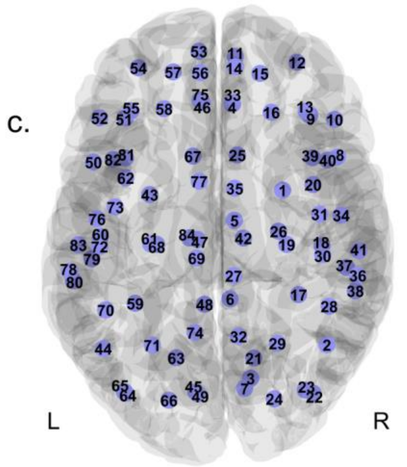


| Right | Left | Abbreviations |
|---|---|---|
| 1 | 43 | AMY |
| 2 | 44 | ANG |
| 3 | 45 | CAL |
| 4 | 46 | ACC |
| 5 | 47 | MCC |
| 6 | 48 | PCC |
| 7 | 49 | CUN |
| 8 | 50 | IFGoper |
| 9 | 51 | IFGorb |
| 10 | 52 | IFGtri |
| 11 | 53 | SFGmedOrb |
| 12 | 54 | MFGorb |
| 13 | 55 | MFG |
| 14 | 56 | SFGmed |
| 15 | 57 | SFGorb |
| 16 | 58 | SFG |
| 17 | 59 | FUSI |
| 18 | 60 | HES |
| 19 | 61 | HIPP |
| 20 | 62 | INS |
| 21 | 63 | LING |

| Right | Left | Abbreviations |
|---|---|---|
| 22 | 64 | IOG |
| 23 | 65 | MOG |
| 24 | 66 | SOG |
| 25 | 67 | SCG |
| 26 | 68 | PHG |
| 27 | 69 | PCL |
| 28 | 70 | IPG |
| 29 | 71 | SPG |
| 30 | 72 | PostCG |
| 31 | 73 | PreCG |
| 32 | 74 | PrCu |
| 33 | 75 | Rec |
| 34 | 76 | ROL |
| 35 | 77 | pSFG |
| 36 | 78 | SMG |
| 37 | 79 | ITG |
| 38 | 80 | MTG |
| 39 | 81 | MTGpole |
| 40 | 82 | STGpole |
| 41 | 83 | STG |
| 42 | 84 | Th |

**S1 Figure. Positions of the 84 nodes in 3D and their abbreviation.**
The nodes are positioned at the centroid of the ROIs of the AAL atlas and represented in 3D along the a.) coronal, b.) sagittal and c.) axial axes. The tables give the abbreviations relative to the different node numbers.
