## Supplementary material for "Phase-amplitude coupling and phase synchronization between medial temporal, frontal and posterior brain regions support episodic autobiographical memory recall": S1 Table

**S1 Table. Anatomical description and abbreviations of the 84 regions used in this study and positions of their centroid in MNI space in mm.**

| Numbers | Abbreviations | Regions | X | Y | Z | Numbers | Abbreviations | Regions | X | Y | Z |
|---|---|---|---|---|---|---|---|---|---|---|---|
| 1 | R AMY | Right amygdala | 26 | 0 | -22 | 43 | L AMY | Left amygdala | -27 | -2 | -22 |
| 2 | R ANG | Right angular gyrus | 44 | -62 | 38 | 44 | L ANG | Left angular gyrus | -46 | -64 | 35 |
| 3 | R CAL | Right calcarine fissure and surrounding cortex | 13 | -76 | 8 | 45 | L CAL | Left calcarine fissure and surrounding cortex | -8 | -82 | 5 |
| 4 | R ACC | Right anterior cingulate cortex | 6 | 35 | 18 | 46 | L ACC | Left anterior cingulate cortex | -6 | 35 | 13 |
| 5 | R MCC | Right median cingulate cortex | 7 | -12 | 40 | 47 | L MCC | Left median cingulate cortex | -8 | -20 | 42 |
| 6 | R PCC | Right posterior cingulate cortex | 5 | -44 | 22 | 48 | L PCC | Left posterior cingulate cortex | -5 | -46 | 25 |
| 7 | R CUN | Right cuneus | 11 | -80 | 28 | 49 | L CUN | Left cuneus | -8 | -82 | 25 |
| 8 | R IFGoper | Right inferior frontal gyrus, opercular part | 48 | 13 | 18 | 50 | L IFGoper | Left inferior frontal gyrus, opercular part | -50 | 11 | 18 |
| 9 | R IFGorb | Right Inferior frontal gyrus, orbital part | 37 | 30 | -16 | 51 | L IFGorb | Left Inferior frontal gyrus, orbital part | -37 | 29 | -15 |
| 10 | R IFGtri | Right inferior frontal gyrus, triangular part | 47 | 28 | 12 | 52 | L IFGtri | Left inferior frontal gyrus, triangular part | -47 | 29 | 10 |
| 11 | R SFGmedOrb | Right superior frontal gyrus, medial orbital | 7 | 53 | -8 | 53 | L SFGmedOrb | Left superior frontal gyrus, medial orbital | -7 | 55 | -7 |
| 12 | R MFGorb | Right middle frontal gyrus, orbital part | 32 | 51 | -12 | 54 | L MFGorb | Left middle frontal gyrus, orbital part | -32 | 49 | -12 |
| 13 | R MFG | Right middle frontal gyrus | 36 | 31 | 33 | 55 | L MFG | Left middle frontal gyrus | -35 | 31 | 34 |
| 14 | R SFGmed | Right superior frontal gyrus, medial | 7 | 49 | 29 | 56 | L SFGmed | Left superior frontal gyrus, medial | -7 | 47 | 30 |
| 15 | R SFGorb | Right superior frontal gyrus, orbital part | 17 | 47 | -17 | 57 | L SFGorb | Left superior frontal gyrus, orbital part | -18 | 47 | -15 |
| 16 | R SFG | Right superior frontal gyrus, dorsolateral | 22 | 31 | 41 | 58 | L SFG | Left superior frontal gyrus, dorsolateral | -21 | 32 | 41 |
| 17 | R FUSI | Right fusiform gyrus | 33 | -42 | -22 | 59 | L FUSI | Left fusiform gyrus | -33 | -46 | -21 |
| 18 | R HES | Right Heschl's gyrus | 42 | -23 | 12 | 60 | L HES | Left Heschl's gyrus | -47 | -19 | 10 |
| 19 | R HIPP | Right hippocampus | 28 | -22 | -12 | 61 | L HIPP | Left hippocampus | -27 | -22 | -12 |
| 20 | R INS | Right insula | 38 | 2 | 1 | 62 | L INS | Left insula | -37 | 5 | 3 |
| 21 | R LING | Right lingual gyrus | 15 | -68 | -5 | 63 | L LING | Left lingual gyrus | -16 | -68 | -5 |
| 22 | R IOG | Right inferior occipital gyrus | 38 | -82 | -10 | 64 | L IOG | Left inferior occipital gyrus | -37 | -81 | -11 |
| 23 | R MOG | Right middle occipital gyrus | 35 | -81 | 20 | 65 | L MOG | Left middle occipital gyrus | -37 | -82 | 15 |
| 24 | R SOG | Right superior occipital gyrus | 23 | -84 | 31 | 66 | L SOG | Left superior occipital gyrus | -19 | -85 | 29 |
| 25 | R SCG | Right subcallosal gyrus | 8 | 14 | -11 | 67 | L SCG | Left subcallosal gyrus | -10 | 13 | -13 |
| 26 | R PHG | Right parahipocampal gyrus | 25 | -17 | -23 | 68 | L PHG | Left parahipocampal gyrus | -24 | -22 | -22 |
| 27 | R PCL | Right paracentral lobule | 7 | -35 | 66 | 69 | L PCL | Left paracentral lobule | -8 | -28 | 67 |
| 28 | R IPG | Right inferior parietal gyrus, excluding supramarginal and angular gyri | 45 | -47 | 50 | 70 | L IPG | Left inferior parietal gyrus, excluding supramarginal and angular gyri | -45 | -48 | 45 |
| 29 | R SPG | Right superior parietal gyrus | 24 | -62 | 59 | 71 | L SPG | Left superior parietal gyrus | -26 | -63 | 56 |
| 30 | R PostCG | Right postcentral gyrus | 42 | -26 | 48 | 72 | L PostCG | Left postcentral gyrus | -47 | -23 | 45 |
| 31 | R PreCG | Right precentral gyrus | 41 | -10 | 49 | 73 | L PreCG | Left precentral gyrus | -41 | -7 | 49 |
| 32 | R PrCu | Right precuneus | 9 | -59 | 41 | 74 | L PrCu | Left precuneus | -9 | -58 | 46 |
| 33 | R Rec | Right gyrus rectus | 6 | 34 | -19 | 75 | L Rec | Left gyrus rectus | -7 | 36 | -19 |
| 34 | R ROL | Right rolandic operculum | 50 | -10 | 14 | 76 | L ROL | Left rolandic operculum | -48 | -12 | 13 |
| 35 | R pSFG | Right posterior superior frontal gyrus | 7 | 0 | 60 | 77 | L pSFG | Left posterior superior frontal gyrus | -7 | 3 | 58 |
| 36 | R SMG | Right supramarginal gyrus | 56 | -35 | 35 | 78 | L SMG | Left supramarginal gyrus | -59 | -33 | 29 |
| 37 | R ITG | Right inferior temporal gyrus | 52 | -31 | -25 | 79 | L ITG | Left inferior temporal gyrus | -50 | -28 | -26 |
| 38 | R MTG | Right middle temporal gyrus | 55 | -41 | -2 | 80 | L MTG | Left middle temporal gyrus | -58 | -37 | -3 |
| 39 | R MTGpole | Right temporal pole: middle temporal gyrus | 38 | 13 | -35 | 81 | L MTGpole | Left temporal pole: middle temporal gyrus | -37 | 13 | -36 |
| 40 | R STGpole | Right temporal pole: superior temporal gyrus | 44 | 12 | -20 | 82 | L STGpole | Left temporal pole: superior temporal gyrus | -42 | 12 | -21 |
| 41 | R STG | Right superior temporal gyrus | 57 | -25 | 7 | 83 | L STG | Left superior temporal gyrus | -56 | -22 | 6 |
| 42 | R Th | Right thalamus | 11 | -20 | 6 | 84 | L Th | Left thalamus | -11 | -20 | 7 |
