## Supplementary material for "Phase-amplitude coupling and phase synchronization between medial temporal, frontal and posterior brain regions support episodic autobiographical memory recall": S2 Table

**S2 Table. Table listing the links of the significant directed network in gamma in the memory condition.**

| Leading | Lagging | Normalized grand average | t value |
|---|---|---|---|
| L Th | R SCG | 1.00 | 4.00 |
| L Th | L AMY | 0.88 | 3.77 |
| L HIPP | L AMY | 0.86 | 4.48 |
| L PCC | L AMY | 0.78 | 5.32 |
| R PCC | R SCG | 0.74 | 3.35 |
| L PCC | L ITG | 0.70 | 3.58 |
| L PHG | L AMY | 0.69 | 3.56 |
| R pSFG | L ACC | 0.61 | 4.98 |
| L pSFG | L ACC | 0.54 | 3.94 |
| R STGpole | L SFGmedOrb | 0.51 | 3.54 |
| R SFGmedOrb | L SFGmedOrb | 0.51 | 4.80 |
| R pSFG | R ACC | 0.47 | 3.60 |
| R SFGmedOrb | L ACC | 0.46 | 3.81 |
| L pSFG | R ACC | 0.43 | 3.33 |
| R pSFG | R SCG | 0.36 | 3.39 |
| L ITG | L MOG | 0.34 | 3.56 |

Annotations: L: left, R: Right, Th: thalamus, SCG: subcallosal gyrus, AMY: amygdala, HIPP: hippocampus, PCC: posterior cingulate cortex, ITG: inferior temporal gyrus, PHG: parahippocampal gyrus, SFG: superior frontal gyrus, pSFG: posterior SFG, ACC: anterior cingulate cortex, STGpole: polar part of the superior temporal gyrus, SFGmedOrb: medial orbital part of the SFG, ITG: inferior temporal gyrus, MOG: middle occipital gyrus.
