## Supplementary material for "Phase-amplitude coupling and phase synchronization between medial temporal, frontal and posterior brain regions support episodic autobiographical memory recall": S3 Table

**S3 Table. Table listing the links of the significant directed network in high theta in the memory condition.**

| Leading | Lagging | Normalized grand average | t value |
|---|---|---|---|
| R PCC | R SCG | 1.00 | 3.74 |
| R PCC | L SCG | 1.00 | 3.90 |
| R PCC | R Rec | 1.00 | 3.67 |
| L PCC | L SCG | 0.94 | 3.75 |
| L PCC | R SCG | 0.93 | 3.61 |
| L PCC | L Rec | 0.91 | 3.47 |
| L PCC | R Rec | 0.91 | 3.42 |
| L PCC | R ACC | 0.84 | 3.29 |
| R PCC | L Th | 0.83 | 3.50 |
| R PCC | R Th | 0.82 | 3.47 |
| L PCC | L Th | 0.79 | 3.42 |
| L PCC | R Th | 0.74 | 3.29 |
| L PCC | R MCC | 0.67 | 3.61 |
| L PCC | L MCC | 0.66 | 3.54 |
| L PrCu | R pSFG | 0.57 | 3.55 |
| R ITG | R SFGorb | 0.52 | 3.99 |
| R ITG | R SCG | 0.51 | 3.47 |
| L PrCu | R MCC | 0.50 | 3.80 |
| R ITG | R SFGmedOrb | 0.44 | 3.30 |
| R ITG | L SFGorb | 0.40 | 3.38 |
| R ITG | R IFGoper | 0.37 | 3.90 |

Annotations: L: left, R: Right, PCC: posterior cingulate corteax, SCG: subcallosal gyrus, Rec: gyrus rectus, ACC: anterior cingulate cortex, Th: thalamus, MCC: median cingulate cortex, PrCu: precuneus, SFG: superior frontal gyrus, pSFG: posterior SFG, ITG: inferior temporal gyrus, SFGorb: orbital part of the SFG, ITG: inferior temporal gyrus, SFGmedOrb: medial orbital part of the SFG.
