## Supplementary material for "Phase-amplitude coupling and phase synchronization between medial temporal, frontal and posterior brain regions support episodic autobiographical memory recall": S4 Table

**S4 Table. Table listing the links of the significant phase-amplitude coupling directed network when contrasting the memory vs. math conditions.**

| Phase | Amplitude | Normalized grand average | t value |
|---|---|---|---|
| R INS | L MTG | 1.00 | 3.93 |
| R ANG | L MTG | 0.95 | 3.51 |
| R MTGpole | L PHG | 0.95 | 3.31 |
| R STG | L FUSI | 0.89 | 3.55 |
| R MTGpole | L Rec | 0.85 | 3.55 |
| R MTGpole | L SFGorb | 0.84 | 3.40 |
| R ANG | L FUSI | 0.82 | 3.43 |
| L HIPP | R SPG | 0.82 | 3.37 |
| L SFGmed | L STG | 0.81 | 4.31 |
| L MCC | R SPG | 0.79 | 3.39 |
| L ITG | L SFGmedOrb | 0.78 | 3.32 |
| L PHG | L SFGmedOrb | 0.77 | 3.36 |
| L HIPP | L SFGmedOrb | 0.77 | 3.35 |
| L STG | L SFGmedOrb | 0.76 | 3.50 |
| R AMY | R SPG | 0.76 | 3.41 |
| L LING | L SFGmedOrb | 0.76 | 3.42 |
| R INS | L Rec | 0.74 | 3.52 |
| R PreCG | L FUSI | 0.71 | 3.83 |
| L MTG | L Rec | 0.64 | 3.42 |
| R SOG | R SPG | 0.61 | 3.52 |

Annotations: L: left, R: Right, INS: insula, MTG: middle temporal gyrus, ANG: angular gyrus, MTGpole: polar part of the MTG, PHG: parahippocampal gyrus, STG: superior temporal gyrus, FUSI: fusiform gyrus, Rec: gyrus rectus, SFG: superior frontal gyrus, SFGorb: orbital part of the SFG, HIPP: hippocampus, SPG: superior parietal gyrus, SFGmed: medial part of the superior frontal gyrus, MCC: median cingulate cortex, ITG: inferior temporal gyrus, SFGmedOrb: medial orbital part of the SFG, AMY: amygdala, LING: lingual gyrus, PreCG: precentral cortex, SOG: superior occipital gyrus.
